# Drivers of genetic differentiation and recent evolutionary history of an Eurasian wild pea

**DOI:** 10.1101/2021.06.14.448334

**Authors:** Timo Hellwig, Shahal Abbo, Ron Ophir

**Affiliations:** The Levi Eshkol School of Agriculture, Hebrew University of Jerusalem, Rehovot 7610001, IL; Volcani Center, Agricultural Research Station, Rishon LeZion, IL

**Keywords:** genetic diversity, genetic differentiation, crop wild relative, range expansion, demographic history

## Abstract

Genetic diversity a major determinant for the capacity of species to persist and adapt to their environments. Unraveling the factors affecting genetic differentiation is crucial to understand how genetic diversity is shaped and species may react to changing environments. We employed genotyping by sequencing to test the influence of climate, space, latitude, altitude and land cover on genetic differentiation in a collection of 81 wild pea samples (*Pisum sativum* ssp. *elatius*) from across its distribution range from western Europe to central Asia. We also attempted to elucidate the species recent evolutionary history and its effect on the current distribution of genetic diversity. Association of single SNPs with climate variables were analyses to test for signatures of local adaptation. Genetic variation was geographically structured into six distinct genetic cluster. Two of which were associated with a taxonomic group (*Pisum sativum* ssp. *humile*) that according to some researchers does not qualify for a sub-species rank due to its alleged lack of genetic distinctness from other conspecific groups. The effect of the tested factors influencing genetic differentiation were rather variable among genetic clusters. The climate predictors were most important in all clusters. Land use was more important in clusters from areas strongly influenced by human land use, especially by agriculture. We found a statistically significant association of 3,623 SNPs (2.4 % of all SNPs) with one of the environmental predictors. Most of them were correlated with latitude followed by temperature, precipitation and altitude. Estimation of SNP effects of the candidates resulted in a missense to silent ratio of 0.45, suggesting many of the observed candidates SNPs may alter the encoded amino acid sequence. Wild peas went through a genetic bottleneck during the last glacial period followed by population recovery. Probably associated with this population recovery, we detected a range expansion, which may have led to an eastward range expansion of the European cluster to Turkey and thereof southwards and eastwards. Overall, the interplay of several environmental factors and the recent evolutionary history affected the distribution of genetic diversity in wild peas where each subpopulations were differently affected by those factors and processes.

## Introduction

Genetic diversity is the raw material of evolution and, therefore, a major determinant of the ability of species to persist and adapt in changing environments (Crozier, 1997; Barret & Schluter, 2008; Prentis et al., 2008; Matuszewski et al., 2015; Lai et al., 2019). It plays a fundamental role in the evolutionary trajectory of species and thus affects all levels of biodiversity (Crutsinger et al., 2006, 2007; Hughes et al., 2008; Stange et al., 2021). This has direct implications for species utilization in agriculture (Kannenberg & Falk, 1995; Hoisington et al., 1999), pharmacology (Elisabetsky & Costa-Campos, 1996; Barata et al., 2016) or bioenergy (Jaradat; 2010; Kiru & Nasenko, 2010) as well as on ecosystems’ stability and scope for recovery (Reusch et al., 2005; Reynolds et al., 2012). Therefore, understanding how species genetic diversity is distributed is crucial. A major determinant of genetic differentiation is the balance between the degree of geneflow between populations and random genetic drift. Geneflow can be restricted due to spatial distribution, if populations are separated by large geographic distances (Slatkin, 1993). A process known as isolation by distance (IBD) that leads to spatial autocorrelation of genetic variation (Wright, 1943). IBD was extensively studied and is widespread in nature (Meirmans, 2012; Sexton et al., 2014). Genetic variation may also be shaped by (expressed as correlation with) environmental variables, which may lead to a pattern of isolation by environment (IBE; Shafer & Wolf, 2013; Wang & Bradburd, 2014). Similarly to the case of IBD, geneflow between populations that inhabit contrasting environments is limited. However, the causes of restriction to geneflow between populations in contrasting environments are more difficult to determine. Several processes like reduced hybrid fitness, genotype-dependent dispersal and natural or sexual selection against immigrants may cause IBE (Servedio, 2004; Nosil et al., 2005; Edelaar et al., 2008; McBride & Singer, 2010; Bolnick & Otto, 2013). IBE has been documented in a variety of different species including barley (*Hordeum vulgare*), soft snow-grass (*Poa hiemata*) or Japanese larch (*Larix kaempferi*) among others (Tanto Hadado et al., 2010; Byars et al., 2009; Nishimura & Setoguchi, 2011; for review see Shafer & Wolf, 2013 and Sexton et al., 2014). A third process, termed isolation by resistance (IBR; McRae, 2006) takes a third dimension of restriction to dispersal and geneflow into account, the heterogeneity of landscapes. The IBR model describes genetic variation in relation to resistance, where the resistance depends on the habitability of landscape features, which connects populations (McRae & Beier, 2007). Cities, cropland or large water bodies present higher resistances to many species compared with forests or grassland and, by that pose larger restrictions on the connectivity of populations. IBR takes this effect into account in addition to the geographic distance and can therefore be considered an extension of the simple IBD model. IBR is particularly important for plants because they cannot easily move through inhabitable areas like cities, cropland or large water bodies and was found to be a better predictor of plants dispersal capabilities than IBD (Cruzan & Hendrikson, 2020, Grasty et al., 2020). Besides those processes that directly affect geneflow patterns among populations, the demographic history of a species can indirectly influence the genetic differentiation between populations. Increased random genetic drift due to a genetic bottleneck may promote the divergence of populations (Wright, 1931). This effect is amplified in peripheral populations, but can also occur in central populations, if the reduction of population size leads to fragmentation, which causes altered geneflow patterns (Lowe et al., 2005; Bouzat et al., 2008; Stevens et al., 2018). An increase in population size can also lead to increased genetic differentiation if accompanied by range expansion. Population expansions may occur in a continuous-gradual manner or via piecemeal punctuations as in a simple stepping-stone model (Excoffier et al., 2009). During a range expansion under the stepping-stone model, the low effective population size of a founding population may lead to increased genetic drift in the population which can drive rapid differentiation between the derived and the source population (Baker & Moeed, 1987). In a series of colonizations, this effect is likely to persist and thereby create an IBD pattern (Ramachandran et al., 2005; Orsini et al., 2013). Naturally, additional parameters will complicate the effects an expansion may have on the involved populations, e.g. the expansion rate or the degree of ongoing geneflow with respective parent populations (Excoffier et al., 2009). Another cause of genetic differentiation following a founder event is described by the monopolization hypothesis (De Meester et al., 2002). After the colonization of a new habitat, rapid adaptation may occur (van Boheemen et al., 2018; Szűcs et al., 2017; Yin et al., 2021) and can provide an adaptive advantage to early founders over late comers leading to evolution-mediated priority effects (De Meester et al., 2016). The resulting adaptive divergence can cause reduced geneflow with the parent population due to the same processes like in the case of IBE (above). This monopolization scenario is equivalent to niche pre-emption (Fukami, 2015), but its priority effects are the result genetic adaptation (De Meester et al., 2016).

The Pea (Pisum L.) genus is commonly divided into two species, *Pisum fulvum* and *Pisum sativum*. *Pisum fulvum* is a rather homogenous entity and a typical east Mediterranean element restricted to Syria, Lebanon, Israel and Jordan where it forms small scattered patches in a variety of habitats (Ben-Ze’ev & Zohary, 1973; Ladizinsky & Abbo, 2015). *Pisum sativum* is divided into at least two subspecies, the domesticated *P. s.* ssp. *sativum* and an aggregate of wild forms collectively named *P. s.* ssp. *elatius* (e.g., Palmer et al., 1985; Hoy et al., 1996; Vershinin et al., 2003; Jing et al., 2007, 2010; Kosterin et al., 2010; Zaytseva et al. 2012; Bogdanova et al., 2018). Older classifications recognized another species, *P. humile* (Boissier, 1867) which was later included within *P. sativum* as a wild subspecies and divided into two varieties, *P. s.* ssp. *humile* var. *syriacum* (often termed northern humile) and *P. s.* ssp. *humile* var. *humile* (known as southern humile, *sensu* Ben-Ze’ev & Zohary, 1973; and see Ladizinsky & Abbo, 2015). Others, however, consider *P. s.* ssp. *humile* a synonym of *P. s.* ssp. *elatius* (Maxted & Ambrose, 2001; Vershinin et al., 2003; Ellis, 2011). Wild *P. sativum* forms (*P. s.* ssp. *elatius* - *sensu lato*) occur across a large geographic range from the Atlantic districts of southern Europe all around the Mediterranean basin, the Black Sea and the Caspian Sea, at least as far east as Turkmenistan (Karakala region) and the Iranian Zagros Mountains (Ladizinsky & Abbo, 2015). Isolated scattered populations also occur as far north as Hungary and Bulgaria (Smykal et al. 2017). Like many Mediterranean elements, *P. s.* ssp. *elatius* exhibits a winter-annual life-cycle, where seeds germinate in autumn after the first precipitation events and overwinter as short statured plants until temperatures rise in the spring. Wild peas are similar to other wild legumes in their strong seed dormancy causing them to build large soil seed banks (Russi et al. 2008; Smýkal et al., 2014; Baskin & Baskin, 2014; Ladizinsky & Abbo, 2015). Genetic diversity in the *P. sativum* complex has been extensively investigated, yet the majority of studies were based on germplasm accessions with a focus on domesticated *P. s.* ssp. *sativum* or rather small sample sizes (Palmer et al., 1985; Hoy et al., 1996; Vershinin et al., 2003; Jing et al., 2007, 2010; Kosterin et al., 2010; Zaytseva et al. 2012; Bogdanova et al., 2018). Exceptions are the studies of Smýkal et al., (2011) and Trněný et al., (2018) which employed a large number of wild samples covering the distribution range of *P. s.* ssp. *elatius*. Yet, neither of them put much emphasis on the factors that may influence genetic variation in wild pea. Smýkal et al., (2018) did investigate the geography and environmental effects on genetic differentiation of wild pea, but only across south-east Turkey.

We attempted to fill the gap left by previous studies and investigated the drivers of genetic differentiation in *P. s.* ssp. *elatius* from a sample set covering its entire distribution range. More specifically, we addressed the following questions: (1) How is genetic diversity in *P. s.* ssp. *elatius* distributed across its native range? (2) What may be the main drivers of genetic differentiation in this taxon? (3) What impact may have the demographic history of this species on its genetic structure? and (4) Can we detect regions of the genome associated with climatic adaptation? Due to its biological characteristics common to many Mediterranean wild legumes and large distribution range covering a great variety of habitats, *P. s.* ssp. *elatius* is a suitable model to study factors driving genetic differentiation that influence the distribution of genetic diversity. Therefore, we consider the present results relevant not only to this species but rather promoting a more general understanding of the factors under which genetic diversity of wild plants evolves.

## Methods

### Plant Material and Genotyping

Our collection contained 81 *P. sativum* ssp. *elatius* individuals which originated from throughout its distribution range (Fig. 1). Some of these accessions were sampled by ourselves, some of them were obtained from genebanks or other sources (Tab. S1). Location details of some genebank accessions were imprecise and appeared in the sea. These were corrected by moving the sample location in a straight line to the adjacent shore (less than ∼2 km in ll cases). Following the classification of Ladizinsky & Abbo (2015), we further subdivided our samples into *P. s.* ssp. *elatius* (48 samples) and *P. s.* ssp. *humile* including its two varieties, *P. s.* ssp. *humile* var. *syriacum* (northern humile; 21 samples) and *P. s.* ssp. *humile* var. *humile* (southern humile; 12 samples) based on morphological characteristics and the respective ecology of the sampling sites. DNA was extracted from the accessions and restriction site associated DNA (RAD) sequencing was conducted by Elshire Group Ltd. (https://www.elshiregroup.co.nz/). Restriction enzyme ApeKI was used for complexity reduction and the protocol of (Elshire et al., 2011) was followed for genotyping by sequencing (GBS). However, 150 bp paired end reads with combinatorial bar codes were produces by the X Ten sequencing platform instead of single end reads with single barcodes by HiSeq 2500 as described in the protocol. Axe-demux (Murray & Borevitz, 2017) was used for demultiplexing. Trimming if adapter and reverse-barcode was done with GBS-PreProcess (https://github.com/Lanilen/GBS-PreProcess; accessed: Feb. 2020). After mapping the processed reads against the *Pisum sativum* ssp. *sativum* reference genome (Kreplak et al., 2019). Stacks (ver. 2.5; Catchen et al., 2011; Catchen et al., 2013) was used to call variants.

**Figure 1:**
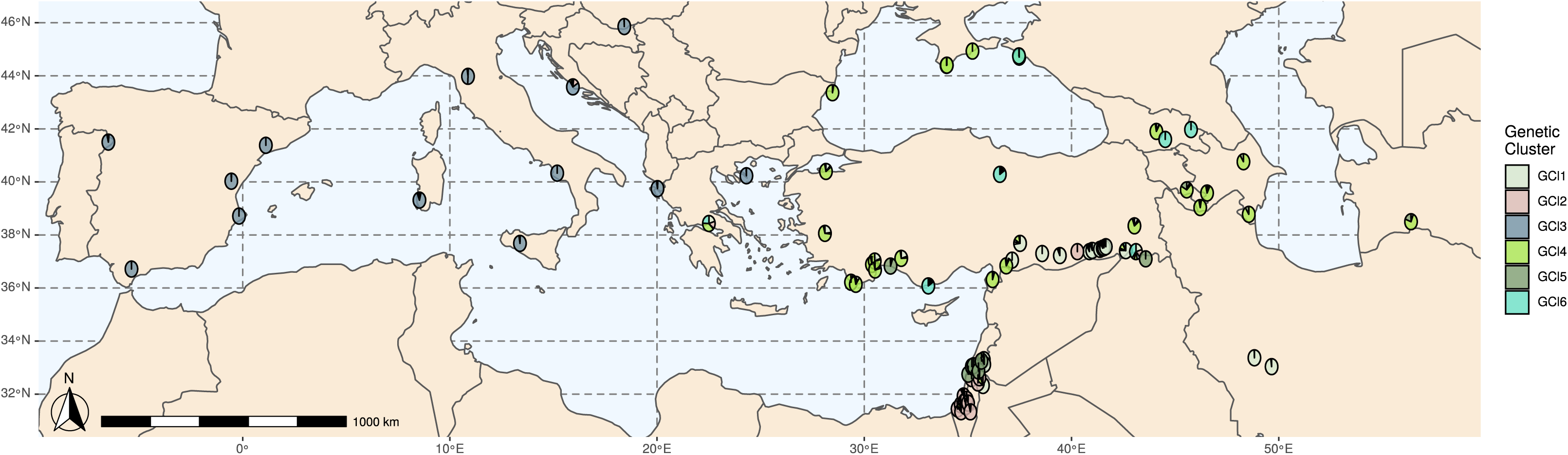
Map of sample locations. Pie charts are located at the sample location of the analyzed accessions and represent sNMF subpopulation fractions with K=6 (for a more detailed map of the Levantinian accessions see Fig. S4).

Only biallelic SNPs were retained. As a minor allele frequency (MAF) threshold we used 0.01. SNPs with less than 40 % genotype calls were removed. We also removed SNPs with minimum depth > 3, minimum mean depth across samples > 5 and minimum genotype quality > 25. The filtering was done with VCFTools (ver. 0.1.17; Danecek et al., 2011). The resulting SNP set was pruned for linkage disequilibrium (LD) with PLINK (v1.90b5.4; Purcell et al., 2007). We shifted 50 kb windows by 5 variants and SNPs were removed when they had r^2^ ≥ 0.5 within one window.

### Population Structure

We used the LD pruned variant set to estimate sample’s subpopulation fractions with sparse non-negative matrix factorization (sNMF; Frichot & François, 2015) in the R statistics environment (R Core Team, 2020) using the package *LEA* (Frichot et al., 2014). We performed sNMF 20 times with numbers of subpopulations (K) ranging from 1 through 20. Since the regularization parameters (ɑ) can lead to biased sNMF results, especially in small data sets, we tested different values to determine an optimum ɑ, which was used in the final analysis. The parameters applied were: 5000 iterations, tolerance = 0.00001, ɑ = 1. For the computation of the cross-entropy criterion, we used 10 % masked genotypes.

We calculated pairwise GENPOFAD distances (Joly et al., 2015) from the LD pruned SNP set to be used in principal coordinate analysis (PCoA). The analysis was done in R with the package *poppr* (Kamvar et al., 2015; R Core Team, 2020).

SplitsTree4 (Huson & Bryant, 2005) was used to create a neighbor-net phylogenetic network based on the entire SNP set. We used the Hasegawa-Kishino-Yano model of nucleotide substitution (Hasegawa et al., 1985). The employed parameters were: transition/transversion ratio = 1.5, empiric base frequencies and a proportion of invariable sites = 0. Since we used a SNP set for the analysis we expected to violate the assumption of equal substitution rates among sites. Therefore, the shape factor of the gamma distribution was set to 0.5 to take wide range of unequal rates among sites into consideration.

We defined a genetic cluster based on subpopulation fractions estimated by sNMF with K=6. An individual was assigned to a genetic cluster when at least 50 % of its subpopulation fraction came from the respective subpopulation (Tab. S1). Two accessions, PeAb15 from Greece and PeAb37 from Turkey, were strongly admixed. They were not assigned to any genetic cluster and excluded from the all subpopulation specific analyses. We calculated pairwise F_st_ between genetic clusters according to (Weir & Cockerham, 1984). Test of significance was done with 9,999 bootstrap values (H_0_: F_st_ = 0). The R package *stampp* (Pembleton et al., 2013; R Core Team, 2020) was used for the computations.

### Landscape genetic analysis

We employed gradient forests analysis, an extension of random forests, to explore non-linear association between climate, land cover, environmental, spatial and allelic variation (Ellis, et al., 2012). The entire analysis was done with the R packages *SoDA* (Chambers, 2013), *vegan* (Jari, 2018), *adespatial* (Dray et al., 2020) and *gradientForest* (Ellis, et al., 2012). The described procedure was done for the entire populations and for each genetic cluster separately.

As environmental variables we used Latitude, Altitude and bioclimatic variables from WorldClim2.1 (Fick & Hijmans, 2017) at 30-seconds spatial resolution. Several of those variables were strongly correlated. There is controversy regarding the question if correlated predictors should be excluded in random forests analyses and if so, where to determine the threshold of correlation (e.g. Cuttler et al., 2012; Murray & Conner, 2009; Knudby et al., 2010). We opted to use an intermediate approach and removed predictors with r² > 0.9 before the analysis. The environmental variables employed in the analysis after removing strongly correlated variables were Latitude, Altitude, BIO11 (mean temperature of coldest quarter), BIO18 (precipitation of warmest quarter), BIO19 (precipitation of coldest quarter), BIO15 (precipitation seasonality), BIO17 (precipitation of driest quarter), BIO10 (mean temperature of warmest quarter), BIO14 (precipitation of driest month) and BIO13 (precipitation of wettest month). In the gradient forest analysis, we used conditional permutation to calculate variables’ importance for predictors with r² > 0.7 to further account for non-independence among predictors (Strobl et al., 2008).

A detailed land cover classification map was obtained from the Copernicus Global Land Service (https://www.copernicus.eu/en; Buchhorn et al., 2019). Due to the high computational demand of the analysis we scaled the resolution down to 1 km. A single resistance value was assigned to each of the 22 land cover classifications (e.g. shrubland: 10, open evergreen needle-leaved forest: 20, cropland: 80, build-up area: 90, sea; 200; Tab. S3). Circuitscape (Circuitscape.jl ver. 5.7.1; Anantharaman et al., 2020) was used to calculated pairwise resistance distances between all samples from the land cover map. In order to use the resistance values in the gradient forest analysis, we transformed the resistance matrix to vectors by performing classical multidimensional scaling (cMDS). We accounted for overfitting by calculating the goodness of fit with varying numbers of retained cMDS coordinates to determine the minimum number of dimensions that retained a sufficient amount of the variation of the original resistance matrix. The cMDS coordinates that still explained 99 % of the variation in the resistance matrix was used as predictors representing land cover in the gradient forests analysis.

Moran Eigenvector Maps (MEMs; Dray et al., 2006) were constructed to model the spatial pattern. For the construction of MEMs, we tested two different algorithms to create the connectivity matrix: Gabriel’s graph and relative neighborhood graph. Each algorithm was tested with three different weights to calculate the spatial weighting matrix (SWM): no weights, linear (1 - D/d_max_, where D was the distance of the node to be weighted and d_max_ the maximum distance between two nodes in the connectivity matrix) and concave down (1 - (D/d_max_)^y^, where y ranged from 2 through 5). The optimum combination of graph and SWM was selected based on the procedure presented by Bauman et al., (2018). The global test of significance was performed with 9,999 permutations. We created a subset of MEMs for each graph - SWM combination using forward selection with double stopping criterion (999 permutations; Blanchet et al., 2008) to avoid overfitting. The graph - SWM combination that showed the highest adjusted R^2^ with the selected MEMs was used for the analysis. Its MEMs represented the spatial pattern in the data.

Gradient forest analysis was implemented in the R package *gradientForest* (Ellis, et al., 2012) where the selected environmental variables, cMDS coordinates and MEMs were used to test their association with the SNP allele frequencies.

### Environmental genome-wide association study

We used redundancy analysis (RDA; Forester et al., 2018) and a latent factor mixed model (LFMM; Caye et al., 2019) to identify SNP associated with environmental predictors. We grouped all temperature and precipitation related variables into the groups temperature and precipitation. Principle component analysis (PCA) was performed on these two groups and the first PCs were used as input for the analsyis. We used Beagle (v5.0; Browning and Browning, 2007; Browning and Browning, 2009; Browning et al., 2018) with 30 iterations, 10 burning steps and effective population size of 50,000 to impute missing genotypes and removed SNPs on scaffolds which left 150,456 SNPs for the analysis. Since we found six well-distinguished populations, we used six latent factors for LFMM analysis. Other input parameters were 100,000 iteration, 10,000 burn-in rounds and ten repetitions per environmental variable. In the RDA, SNPs were considered as candidates if their loadings in the ordination space deviated more than three standard deviations from the mean (approximately equivalent to a two-tailed p-value of 0.0027). Note, that this threshold is rather low, but after the analyses, we only considered SNPs that appeared to be outliers in both analyses as candidates. This approach has been shown to produce rather conservative results and to be a good procedure for a first exploratory analyses of loci under selection (Forester et al., 2018). The analysis were done using ‘LEA’ and ‘vegan’ R packages (Frichot and François, 2015; Jari, 2018). We used the program SnpEff (v. 5.0) to predict effects of candidate SNPs (Cingolani, 2012).

### Statistics of diversity and neutrality

Several diversity statistics were calculated for each of the genetic cluster. Nei’s gene diversity (H_exp_; Nei, 1978), observed heterozygosity (H_obs_) and nucleotide diversity (π) were calculated with *adegenet* (Jombart & Ahmed, 2011) *and VCFTools* (ver. 0.1.17; Danecek et al., 2011). Haplotypes were generated by HaploView (Barrett et al., 2005) using the four gamete rule (Wang et al., 2002). Markers which heavily violated HWE were removed from the analysis to ensure that SNPs under selection are not included in the analysis. We applied Bonferroni correction to account for multiple testing and removed markers with a p-value below 0.05/(number of markers). We calculated number of haplotypes (Ht_n_) and haplotype diversity (Ht_d_) from the results of HaploView using R core functions (R Core Team, 2020).

Tests of neutrality were conducted by calculating Tajima’s D (Tajima, 1989) and R_2_ (Ramos-Onsins & Rozas, 2002) for each genetic cluster using the R packages *r2vcftools* and *pegas (Pope, 2020; Paradis, 2010).* Significance of both statistics were accessed with simulations of 100,000 neutral sequences.

We estimated selfing rates of the entire population using the method of David et al. (2007). P-values of the the identity disequilibrium g_2_ were calcualted with 9,999 permutations.

### Inference of demographic history

Inference of demographic history was done using Stairway Plot 2 (Liu & Fu, 2020). The analysis was run with the LD pruned variant set and 1,000 replicates to estimate confidence intervals. Although wild peas are annual plants, their generation time is longer than one year due to their strong seed coat imposed physical dormancy. Based on observations in our experimental farm where volunteers emerged up to six years after growing wild peas, we opted to use an average generation time of three years. Mutation rates in peas are unknown. In line with the generation time hypothesis, which states that species tend to have accelerated rates of molecular evolution if they have shorter generation times, we used a mutation rate of 2e-9, since this is at the lower end of the estimated mutation rates of *Arabidopsis thaliana* (Exposito-Alonso et al., 2018; Lynch, 2010). This reflects the higher generation time that we used compared to the 1-year generation time of *A. thaliana*. We used the unfolded sites frequency spectrum (SFS) as input for Stairway Plot 2 and ignored singletons as suggested by the authors (Liu and Fu, 2015).

### Detection of range expansion

We tested for a past range expansion using the directionality index (ψ) (Peter and Slatkin, 2013; Peter and Slatkin, 2015). Since this approach assumes neutrality, we excluded SNPs that violate HWE using the same approach as for the calculation of neutrality statistics (see above). We additionally removed sites with more than 10 % missing genotypes to ensure that ψ, which detects clines in allele frequencies, can be calculated reliably. The null distribution of the ψ was generated by permuting the allele frequencies of each SNP and assigning them randomly to a population. P-values were calculated based on this null distribution (H_0_: ψ = 0). Since permutations tend to underestimate the variance in the data if SNPs are in linkage disequilibrium (Peter & Slatkin, 2013), we used the LD pruned variant set for the analysis.

## Results

### Genotyping by sequencing

We detected 1,973,460 variants in total while 167,766 SNPs were retained after quality filtering and 106,770 after LD pruning. In the entire SNP set as well as in the LD pruned set, SNPs were distributed throughout the entire genome without notable large gaps (Fig. S1). In both SNP sets the majority of sites had low missing genotype calls with a median and mean of 11 % and 14 %, respectively (Tab. S2). Rare alleles were abundant in both SNP sets. The third quantile of MAF was 0.13 and 0.11 in the filtered and the LD pruned set, respectively (Tab. S2). Observed heterozygosity was extremely low with only 2 % at the third quantile and a mean of 0.01 (Tab. S2). In the entire variant set the mean and median of the nucleotide diversity (rr) were 0.09 and 0.16, respectively and ranged from 0.04 in the first quantile to 0.233 in the third quantile. After LD pruning similar values were observed (Tab. S2).

### Population Structure

Analysis of population structure exposed six well distinct clusters (Cl1 - Cl6; Fig. 2A) with F_st_ values ranging from 0.162 (Cl1 - Cl4) to 0.438 between Cl3 and Cl6 (Tab. 1). The majority of individuals only showed small degrees of admixture in the sNMF analysis and only two accessions (PeAb15 and PeAb37) did not have a subpopulation fraction of > 0.5 of any genetic cluster and were not assigned to any cluster (Fig. 2A). Individuals from Cl1 were sampled from the northern and eastern part of the Fertile Crescent. Cluster 4 and Cl6 showed a wide distribution range from Turkey to the Caucasus and around the Black Sea while Cl2 and Cl5 were located in geographically restricted areas in southern and northern Israel, respectively. The sample locations of Cl3 were spread all over southern Europe (Fig. 1). Cluster 1 and Cl4 split from a common node in the phylogenetic network and their individuals cluster closely together in PCoA (Fig. 1 B,C). They also cluster together until K = 4 in the sNMF analysis (Fig. 2A). Cl2 and Cl5 also shared the same origin in the phylogenetic network. Interestingly, Cl3 emerged from the same node, although it was located at the opposite end of the distribution range of the germplasm collection relative to Cl2 and Cl5. PCoA results were in line with these observations (Fig. 1 B, C). Cluster 6 was placed between the two main branches. Despite the apparent geographic pattern, some outliers were observed. One sample from Cl5 was sampled from eastern and one from western Turkey. The sample location of two individuals assigned to Cl1 were located in Jordan and on the Golan Heights close to the Israeli - Syrian border.

**Table 1:**
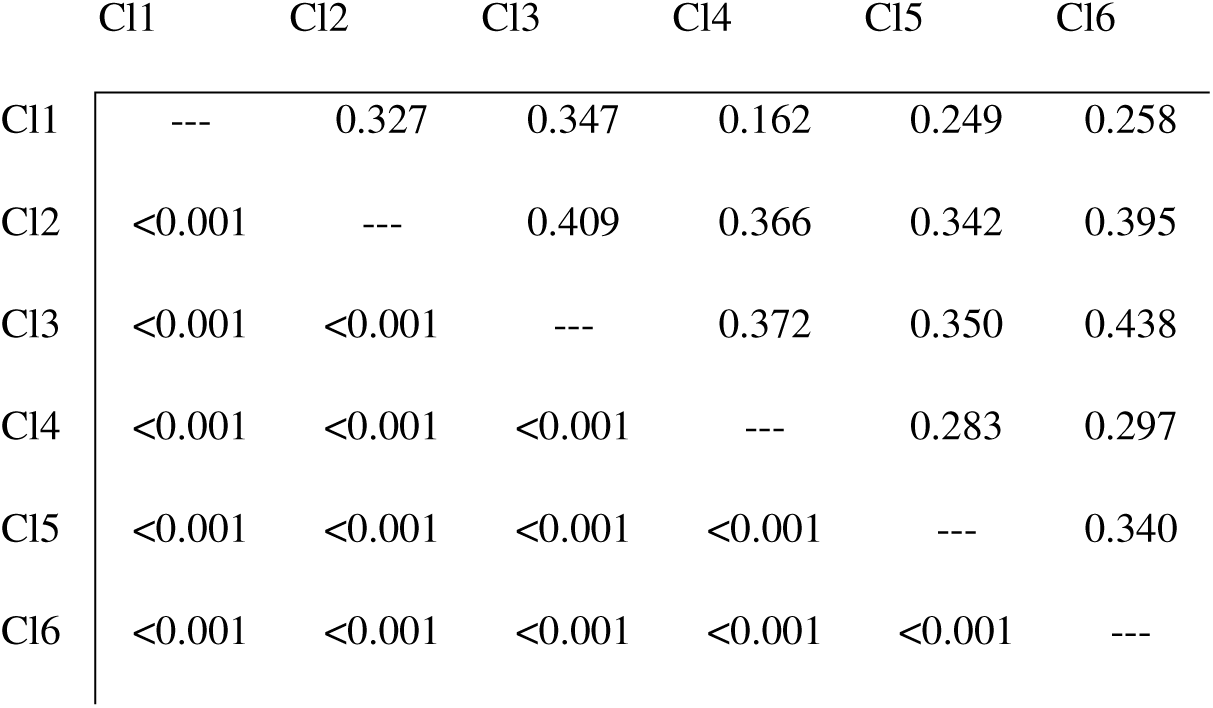
Pairwise F_st_ values between genetic clusters (upper triangle) with corresponding p-values based on 999 bootstrap values (lower triangle).

**Figure 2:**
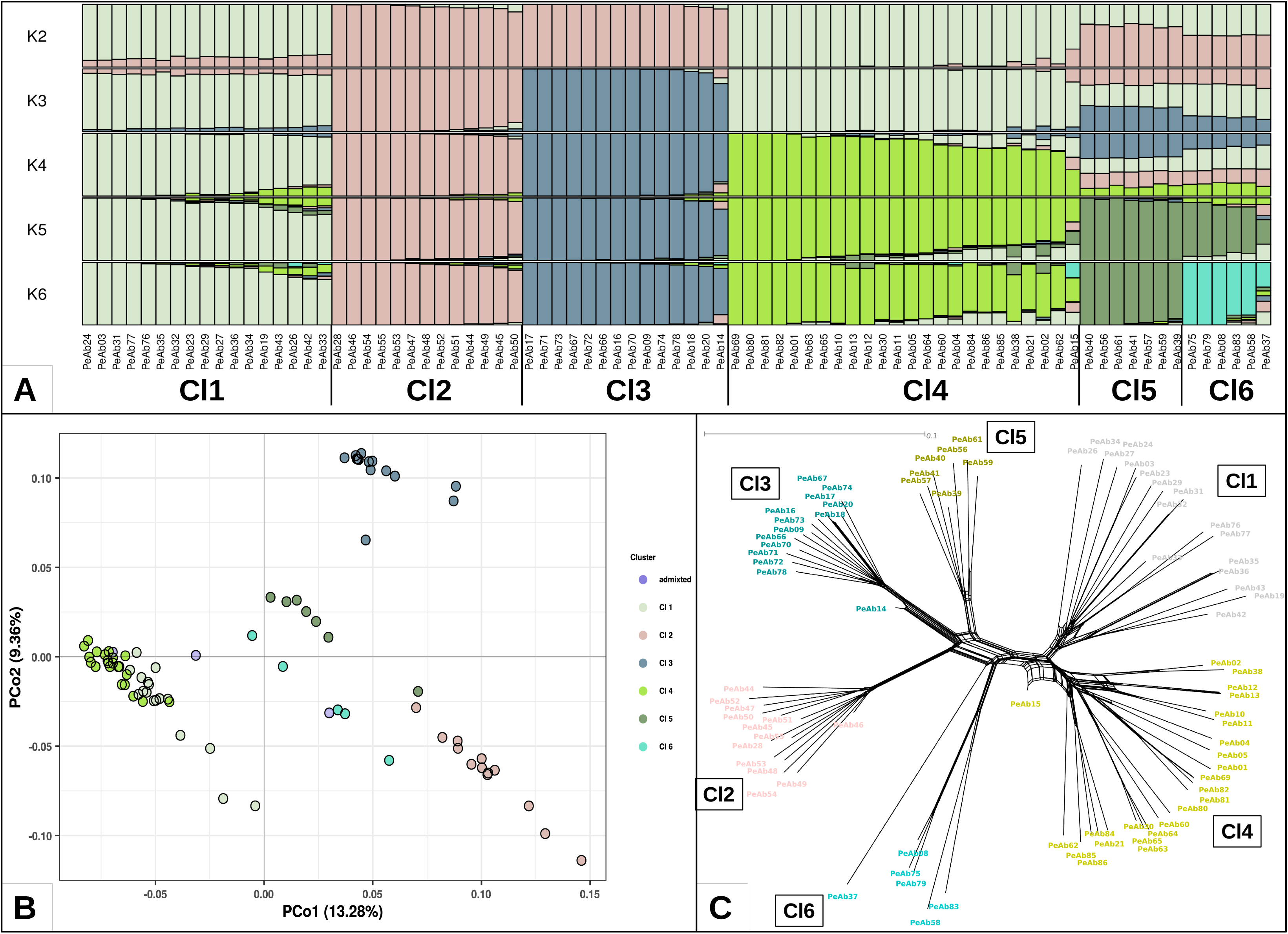
Results of population structure analysis. A: subpopulation fraction estimated by sNMF with K ranging from 2 through 6. B: PcoA; percentages in axes titles represent the fraction of explained variation; accessions that did not have a subpopulation fraction of > 0.5 of any genetic cluster were defined as admixed. C: Neighbor-net phylogenetic network.

### Landscape genetic analysis

In the gradient forest analysis 21.00 % of the genetic variation could be explained by the employed variables, where the spatial predictors were most influential (10.60 %) followed by the climate (7.28 %; Fig. 3). Within the climate predictors precipitation related variables had higher importance relative to temperature related variables (Fig. S2). Altitude, land cover and latitude accounted for minute fractions of the genetic variation with 0.47 %, 1.20 % and 1.45 %, respectively. In the individual genetic clusters the effect of the different predictors were overall similar with few exceptions. Notably, small association was estimated for Cl6 with only 1.06 % in total. The very small sample size of this cluster has to be kept in mind and its estimate should be considered with caution. Climate showed the strongest association with genetic variation in all clusters, altitude the weakest. Within the climate predictors, precipitation related variables had an overall stronger effect than temperature related variables (Fig. S2). The effect of land cover was particularly strong in Cl1 (9.55 %) and Cl2 (6.17 %), where it was the second strongest effect after climate and followed by the geography. The spatial predictors showed strong association in Cl1, Cl3 and Cl4, which were also the clusters with the widest distribution range.

**Figure 3:**
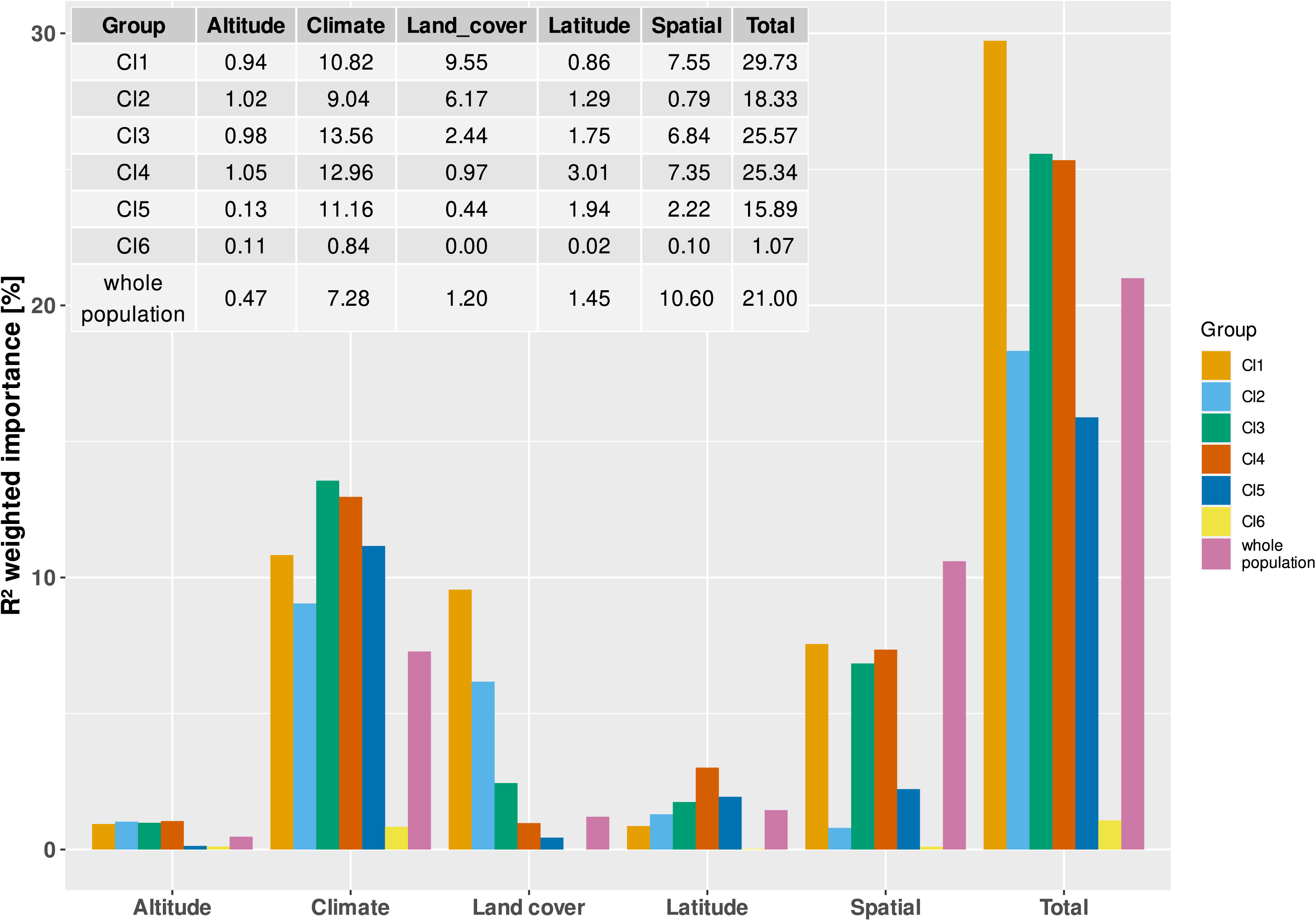
R² weighted importance of environmental, spatial and land cover variables for each genetic cluster and the whole population from gradient forest analysis (in percentages).

### Environmental genome-wide association study

We detected 3,623 SNPs (2.4 % of all SNPs) that were significantly correlated with one of the environmental predictors. Most SNPs were associated with latitude (1,140) followed by temperature (963), precipitation (929) and altitude (591). Candidate SNPs were distributed rather evenly across the genome with gaps in centromeric regions. Notably, only very few of the SNPs with the highest correlation were located on chromosomes six and seven (Fig. S3). The ratio of missense to silent SNPs was 0.45. Intron variants were the most abundant predicted effect (29.9 %) followed by synonymous SNPs (27.4 %). Thirty seven SNPs possibly caused a premature start codon and one SNP created a stop codon within a gene. All candidate SNPs, their predicted effects and the corresponding genes are depicted in Tab. S4.

### Statistics of diversity and neutrality

Observed heterozygosity was very small and barely varied across genetic clusters (Tab. 2). Expected heterozygousity was also small and only little variation was observed among clusters ranging from 0.07 in Cl3 to 0.1 in Cl1 (Tab. 2). Nucleotide diversity was highest in Cl5 and Cl6 with 0.38 and 0.46, respectively. These two clusters also had the highest Ht_d_ (both 0.92) and the fewest number of haplotypes with 3,811 and 2,893, respectively (Tab. 2). The lowest π (0.24) and Ht_d_ (0.74) was found in Cl4 which also had the lowest Ht_n_ (4045) after Cl4 and Cl5. Cluster 1, Cl2 and Cl3 showed intermediate values of π, Ht_d_, Ht_n_ (Tab. 2). Cluster 4 had a low negative Tajima’s D value (−0.12; p < 1e-05). Clusters 1 to 3 had very low positive Tajima’s D values, yet all of them were significantly different from 0. Tajima’s D was high for Cl5 and Cl6 with 0.58 (p < 1e-05) and 0.89 (p < 1e-05), respectively. Contrary to the Tajima’s D values none of the R_2_ values were significant even though they ranged from 0.12 in Cl1 to 0.2 in Cl6 (Tab. 2). We estimated a selfing rate of 0.700 for the total population

**Table 2:**
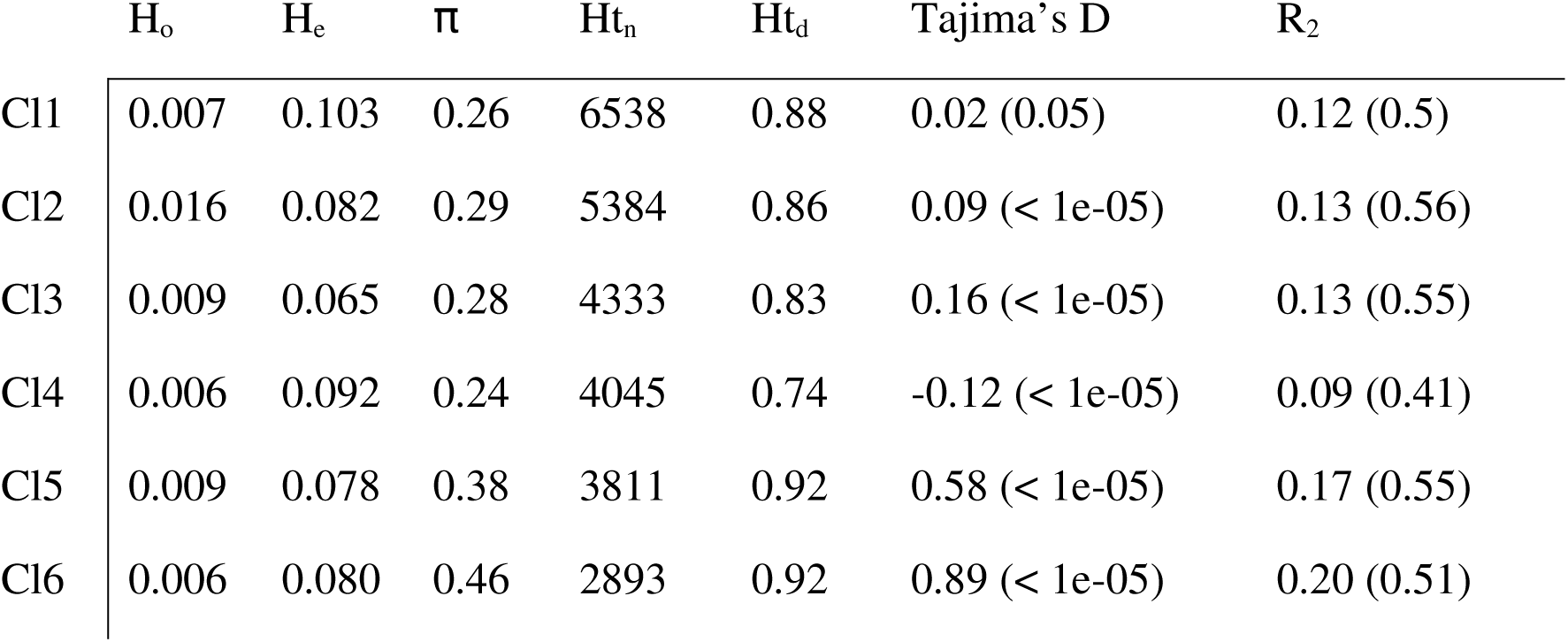
Statistics of diversity (H_o_: observed heterozygosity, H_e_: expected heterozygosity, *π* : nucleotide diversity, Ht_n_: number of haplotypes, Ht_d_: haplotype diversity) and neutrality of each genetic cluster (Tajima’s D and R_2_).

### Inference of demographic history

Estimations of past effective population sizes (N_e_) depicted an increase of N_e_ during the early phase of the last glacial period from around 150 k to 90 k year ago. Afterwards, a genetic bottleneck slowly started and became increasingly rapid until N_e_ dropped from ∼180 k to 60k roughly 3 k years ago. The population quickly recovered after the bottleneck and depicted constant N_e_ on a similar level as before the bottleneck for the last 1.5 k year. Note that we refrained from depicting results deeper into the past beyond around 100 - 200 thousand years, because estimations tend to become less reliable for such periods. This is especially true for results of Stairway Plot 2 (Liu & Fu, 2015).

### Detection of range expansion

Significant directionality indices (ψ) were detected for Cl3 indicating an expansion towards Cl6 and Cl1 which were also the highest ψ values with 0.32 and 0.30, respectively. Both, Cl4 and Cl5 had significant ψ leading to Cl1 and Cl2, but ψ values originating from Cl4 were higher than those of Cl5 (Tab. S5). Overall, we detected expansion from the eastern part of the distribution area to the central part and from there southwards and further to the east (Fig. 5).

## Discussion

### Structure of genetic diversity

We observed six well distinct genetic clusters showing a hierarchical structure with a notable geographic pattern (Tab. 1, Fig. 1, 2). Overall our genetic structure is congruent with the structure reported by Smýkal et al., (2017). Large overlap could be observed between Cl3, Cl4, and Cl1 with the subpopulations Q3, Q4, and Q5 reported in Smýkal et al. (2017), respectively. These are also largely equivalent to the clusters 9, 10 and 1 of Trněný et al. (2018) who used the same wild germplasm array of Smýkal et al., (2017) but additionally included the two domesticated taxa. Yet, if viewed in greater detail, there are some differences, e.g. Smýkal et al., (2017) and Trněný et al. (2018) could not detect distinct subpopulations in the southern Levant, most likely due to the smaller number of samples from this area. Therefore, they could not have observe that Cl2, Cl3 and Cl5, which were sampled from Israel (Cl2, Cl5) and southern Europe (Cl3), showed close genetic proximity even though they are geographically remote. This pattern was also observed by Kosterin et al. (2010) who investigated the phylogeny of wild peas based on three dimorphic markers (nuclear, plastid and mitochondrial) and divided the genus into four lineages (A through D). They suggested that lineage A represents the ancestral state since its marker combination is present also in *P. fulvum*. Combination A is also dominant in *P. s.* ssp. *elatius* samples from Israel and the Mediterranean islands Sardinia and Menorca while lineage C, which is distinct from A by only a single mutation, could be found in continental southern Europe (France, Greece, Italy). These authors proposed a westward expansion during the Pleistocene climate cooling when oscillating sea levels temporally connected certain Mediterranean islands. Analysis of spatial diversity of niche patterns in wild peas suggested that the center of diversity is currently located in the Near East (Smýkal et al., 2017) which is also supported by our findings as evident from the diverse population structure and the higher diversity statistics of the Near Eastern clusters compared to the European Cl3 (Tab. 2). The same analysis predicted a center of high diversity in northern Africa and the authors proposed that the route through north Africa might be a so far overlooked route during the westward expansion of wild pea. Both proposed scenarios (Kosterin et al. 2010; Smýkal et al., 2017) are consistent with our findings of close genetic proximity of the southern Levantine clusters (Cl2, Cl5) with the southern European Cl3. Unfortunately, our sample collection did not contain any samples from north Africa and the number of available genebank accessions from north Africa is very limited and often with questionable passport data. Sample expeditions to this area may enable researchers to shed more light on the role of this area in past expansion of wild pea. Indeed, the germplasm collection of Smýkal et al., (2017) held a small number of genebank accessions from north Africa, yet the study was criticized for data confusion, including a misclassification of at least one domesticated sample as wild which suggests that one should view parts of this collection with some caution (Kosterin & Bogdanova, 2018).

For decades botanists followed the classification of Boissier (1867) who recognized three wild pea species, *P. fulvum*, *P. elatius* and *P. humile*. Ben-Ze’ev & Zohary (1973) showed by crossing experiments that although chromosomal interchanges distinguish between *P. s.* ssp*. elatius* and the so called northern *P. humile*, *P. s.* ssp*. elatius* and the southern *P. humile* share the same cytotype. Therefore, Ben-Ze’ev & Zohary (1973) considered *P. elatius* and *P. humile* as interfertile (and likewise with the domesticated *P. s.* ssp. *sativum*) and consequently, adopted the classification proposed by Davis (1970), who recognized only two *Pisum* species, *P. fulvum* and the *P. sativum* complex (with its several subspecies), which is mostly accepted today. More detailed classifications within the *P. sativum* complex were proposed in a number of studies, which led to a confusing number of proposed epithets and taxonomic entities that often were considered synonymous with each other (e.g., Maxted & Ambrose, 2001; Ellis, 2011). A unified taxonomic classification of the *P. sativum* complex is not yet agreed upon, but most researchers today follow the classification of Maxted & Ambrose (2001), who divided the *P. sativum* complex into a domesticated subspecies, *P. s.* ssp. *sativum*, and a single wild subspecies, *P. s.* ssp. *elatius*. Consequently, *Pisum s.* ssp. *humile* is nowadays widely considered a synonym of *P. s.* ssp. *elatius*, because it was argued that it is not genetically distinct from studied *P. s.* ssp. *elatius* accessions (Vershinin et al., 2003; Ellis, 2011). However, these studies included only a single *P. s.* ssp. *humile* accession. This may be due to lack of genuine wild samples or due to the fact that the different wild *P. sativum* subgroups (in our view, ecotypes) are often difficult to distinguish morphologically. This may be problematic when accessions are grown in greenhouses, outside of their natural ecological zone, by researchers who never observed them in their native habitats or in common garden experiments when these stocks maintain their distinct phenotypes. For an example of the profound effect of growth conditions on the morphology see comments and photos in Kosterin et al., (2020). Besides *P. s.* ssp. *elatius*, we included *P. s.* ssp. *humile* and its proposed separation into two varieties, *P. s.* ssp. *humile* var. *syriacum* (northern humile; 21 samples) and *P. s.* ssp. *humile* var. *humile* (southern humile; 12 samples) following Ladizinsky & Abbo (2015). Out of 21 northern *humile* samples 17 clustered in Cl1, which also included the *P. s.* ssp. *elatius* accession PeAb03 (Fig. 2; Tab. S1). All samples classified as southern humile clustered in Cl2, which contained only another sample that was classified as northern *humile* (PeAb28). This contradicts the results of Vershinin et al. (2003) and Ellis (2011) and suggests that northern and southern *humile* are distinct groups within the *P. sativum* complex. Yet, they are just as distinct from each other as they are from other *P. s.* ssp. *elatius* groups. It should also be noted that according to our results *P. s.* ssp. *humile* does not take up a special place in the *P. sativum* complex. From a genetic standpoint, the *P. s.* ssp. *humile* groups are just two out of six well distinct entities within this complex.

### Drivers of genetic diversity

We observed notable variation among genetic differentiation explained by the employed variables between clusters. Yet, some common patterns could be detected. In total, all variables accounted for a comparable fraction of genetic variation (Fig. 3). Cluster 6 was an outlier with a very small fraction of genetic variation associated with the employed predictors. As mentioned above, the small sample size of this cluster (5) could have biased the analysis and the results of this cluster should be viewed with caution.

The bioclimate appeared to be the strongest driver of genetic differentiation in *P. s.* ssp. *elatius* since the bioclimate variables were most strongly associated with genetic structure in all genetic clusters and accounted on average for 57 % of the explained variation. Overall, precipitation related variables had a stronger effect than temperature related variables (Fig. S2). These findings differ from the results of Smýkal et al. (2018) who did not find any signal of IBE in *P. s.* ssp. *elatius* populations from south-east Turkey. The limited geographic scale and lower environmental heterogeneity may be the reason why Smýkal et al. (2018) did not detect IBE. Limited IBE was also reported for *P. fulvum* from the southern Levant in a rather small study area, yet with great environmental heterogeneity. The authors attributed the absence of a strong IBE to strong genetic bottleneck in *P. fulvum*, which may have caused random genetic drift to obscure the signal of genome-wide differentiation following the environmental cline (Hellwig et al., 2020, 2021). Isolation by environment is very common in nature, for example, in a meta-study 74 % (52 out of 70) of phylogeographic studies reported patterns of IBE (Sexton et al., 2013). The average effect size of IBE was estimated to be 0.34 when controlled for spatial autocorrelation (Shafer & Wolf, 2013). Our estimated effect was significantly higher which emphasizes the importance of the climate on genetic differentiation in *P. s.* ssp. *elatius.* These findings are particularly important for the utilization of *P. s.* ssp. *elatius* for breeding efforts, since it suggests a large fraction of adaptive variation is related to climatic factors. We found 963 (0.64 % of total) and 926 (0.62 % of total) SNPs significantly correlated with temperature and precipitation, respectively (Tab. S4; Fig. S3). A large fraction of these were non-synonymous polymorphisms within predicted genes (Tab. S4). If these candidates can be validated, they may be useful for crop improvement, because *P. s.* ssp. *elatius* is interfertile with the domesticated *P. s.* ssp. *sativum*.

Although most of the detected candidate SNPs putatively under selection were associated with latitude, the genome-wide effect of latitude was comparably small. These were rather unexpected results, especially for the entire population as well as for clusters with a large north-south distribution like Cl4 and Cl3. It is expected that a number of developmental traits change with latitude as an adaptation to varying day length. This has been observed in a variety of plant species like *Arabidopsis* (Riikimäli et al., 2005; Sami et al., 2008; Fabian et al., 2012; Debieu et al., 2013), soybean (Lu et al., 2015), wild lupin (Mousavi-Derazmahalleh et al., 2018), wild wheat (Matsuoka et al., 2008) and a number of different forest herbs (De Frenne et al., 2009). Local adaptation without genome-wide differentiation following the respective environmental cline has also been observed in *P. fulvum* (Hellwig et al., 2021) and likewise with lava flow lizards (Krohn et al., 2019). Krohn et al. (2019) argued that ongoing migration between populations may counteract genome-wide divergence. It has also been pointed out that genome-wide differentiation is less likely to occur if adaptive loci are not linked to traits involved in reproductive isolation (Slatkin, 1987; Presgraves, 2013). Yet, extensive geneflow over thousands of kilometers across the entire north-south distribution range of *P. s.* ssp. *elatius* is highly unlikely and loci involved in adaptation to latitude (i.e. growth temperature, day length) are likely to be linked to reproductive traits like flowering time. Hellwig et al. (2021) explained their observed pattern with the genetic bottleneck underwent by the studied populations. They hypothesize that neutral areas of the genome are strongly influenced by random genetic drift caused by the reduced N_e_ while adaptive areas are shaped by selection. Even though we also observed a genetic bottleneck during the last glacial period, the scenario hypothesized for *P. fulvum* cannot explain the pattern observed for latitude, because we would expect to see a similar pattern for the other predictors. The age of the adaptation to latitude could be a possible explanation. With time, the linkage between adaptive and the surrounding neutral loci may break apart, leaving adaptive SNPs without associated genome-wide differentiation. However, we do not have any evidence to support this scenario, which therefore remains speculative.

The spatial predictors were most important in clusters with large geographic range. Euclidean distances between sample locations are not considered as meaningful predictor for genetic differentiation since it cannot account for landscape heterogeneity (Slatkin, 1985; McRae, 2005). Indeed, a straight line between sample locations is not a good approximation of the connectivity of plant populations. Despite this criticism, it is still commonly included in landscape genetics analyses. Therefore, we included absolute distance as a predictor in our analysis to make our results comparable with other studies, even though we agree with the criticism and advocate to use more meaningful variables to model the spatial connectivity between populations (e.g. resistance maps like used herein or least cost paths; for review see Zeller et al., 2012).

Land cover had a particularly large impact on genetic differentiation in central Israel (Cl2) and south-east Turkey (Cl1). Central Israel is highly populated and the areas suitable for wild peas is heavily fragmented by roads, industrial estates, urban areas and cropland. Similarly, there are large blocks of arable land in southern Turkey. Land cover was also an important predictor for the distribution of genetic variation in south Europe (Cl3), yet, to a smaller degree relative to the Levant. Southern Europe is also strongly impacted by agriculture, but contrary to the Levant, there are large areas covered by forests, herbaceous vegetation and shrubland that are overall rather well connected. These areas may serve as routes that connect *P. s.* ssp. *elatius* populations and can reduce the effect of habitat fragmentation. Habitat fragmentation is certainly an important process affecting genetic diversity. Habitat fragmentation reduces N*e* which negatively affects genetic diversity due to increased genetic drift (Ellstrand & Elam, 1993; Young et al., 1996). Honnay & Jacquemyn (2007) supported this theoretical prediction in a meta-study and emphasized that such a population genetic response to habitat fragmentation can be observed in rare as well as common species. Another meta-study demonstrated that regardless of the ecological or life-history traits of the investigated species, fragmentation is likely to negatively impact plant sexual reproduction (Aguilar et al., 2006). However, not all species react in the same way to habitat fragmentation. Encinas-Viso et al. (2020) showed that reduced forested area had a significant impact on the genetic differentiation of the Australian legume *Acacia salicina* but not on *A. stenophylla*, which inhabits largely the same area. Foré et al. (1992) even found decreased genetic differentiation in post-fragmentation populations compared to pre-fragmentation populations in *Acer saccharum* from the USA. The results of Prober & Brown (1994), who studied the effect of habitat fragmentation, due to cropland expansion, on *Eucalyptus albens*, indicate that there may be a fragmentation threshold and it is likely that the duration and intensity of habitat fragmentation plays an important role in the plant population genetic response. Our results suggest that one possible explanation is that habitat fragmentation due to intensive human land use, especially agriculture, causes increased genetic differentiation in wild pea. Yet, we cannot say which of the possible responses to habitat fragmentation have caused this increased genetic differentiation.

### Demographic history

Our results suggested that *P. s.* ssp. *elatius* went through a genetic bottleneck during the last glacial period (Fig. 4). A similar population contraction was reported for the wild pea species *P. fulvum* (Hellwig et al., 2020, 2021). Yet, contrary to the situation in *P. fulvum*, N_e_ of *P. s.* ssp. *elatius* increased shortly after the bottleneck and post-recovery estimates of N_e_ after the recovery were even slightly higher than before the contraction. Hellwig et al., (2020, 2021) mentioned several possible reasons why pea populations may have decreased during the last ice age like sea level fluctuations or the increased area covered by forests which is a less suitable habitat for peas. The bottleneck continues into the early phase of the Holocene when agricultural practices were already widespread within our study area and therefore, human made environmental disturbances like deforestation and increased grazing pressure by livestock likely affected pea populations as well. However, this cannot explain why *P. s.* ssp. *elatius* recovered from the bottleneck while *P. fulvum* did not. The fairly similar ecological affinities (including spatial distribution patterns) of the two wild pea species adds to this riddle. Our estimate of the overall selfing rate of *P. s.* ssp. *elatius* (0.700) is higher than the reported estimate of *P. fulvum* (0.495; Hellwig et al., 2021). Populations can benefit from higher selfing rates after a bottleneck, because it ensures reproduction in absence of mating partners (Cheptou, 2019). Therefore, the difference in reproduction pattern between *P. sativum* and *P. fulvum* may be considered as one of the factors that contributed to the difference in the recovery from the genetic bottleneck.

**Figure 4:**
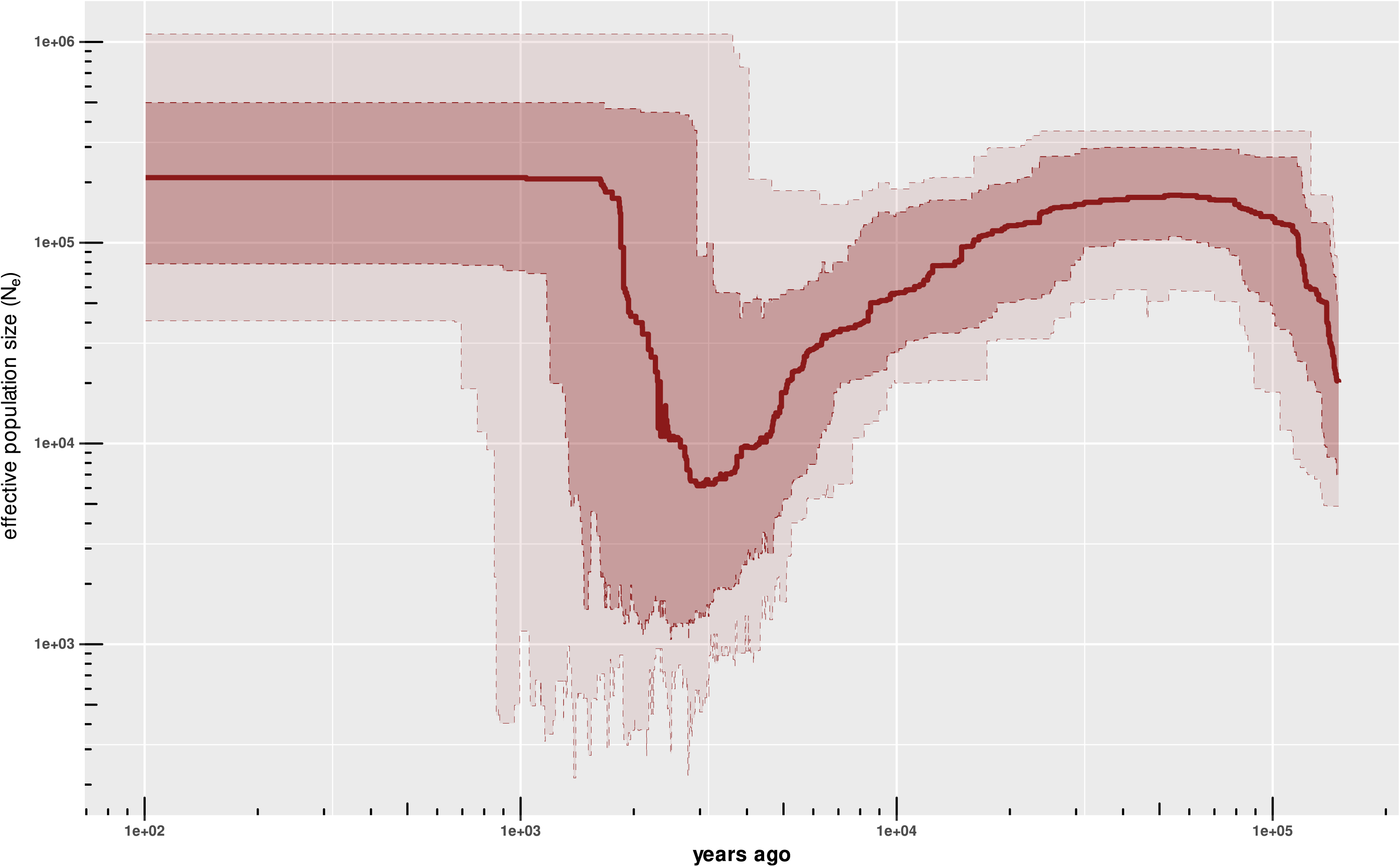
Inferences of past effective population size. The solid line represents the median N_e_. Red and pink shaded areas represent the 97.5 and 87.5 percentiles, respectively. Both axes are on log _10_ scale.

It is expected that a range expansion will create a gradient of genetic diversity along the expansion route due to serial founder effects where variation decreases with increasing distance from the origin (Austerlitz et al., 1997; Edmonds et al., 2004; Ramachandran et al., 2005; Slatkin & Excoffier, 2012). We did not observe such a gradient of genetic variation in our data and statistics of neutrality also did not support a recent population expansion (Tab. 2), although the estimation of historic N_e_ resulted in a population recovery after a strong genetic bottleneck during the last glacial period (Fig. 4). Cluster 4 had a significantly negative Tajima‘s D value indicating a possible population expansion after a bottleneck. However, this value was rather low and, hence, this expansion was likely not severe enough to explain the detected increase of N_e_. In the other clusters, values of Tajima‘s D were significantly positive, which rather speak for a recent population contraction or alternatively, for balancing selection. Yet, Tajima‘s D in those clusters was very close to 0 except for Cl5 and Cl6, where the low sample size may have biased the results. Estimates of R_2_ were not significantly different from 0 in any cluster contradicting the calucaltions of Tajima‘s D. The results of summary statistics are ambiguous and do not allow any clear conclusions. Another population genetic signal of recent range expansion are clines in allele frequencies along the expansion trajectory (Slatkin & Excoffier, 2012). The results of the approach of Peter & Slatkin (2013, 2015) suggest a range expansion rejecting the null model of isolation-by-distance at equilibrium.

Note, that while the center of diversity (hence, the likely origin) of the genus as a whole is in the east Mediterranean basin, the detected clines of allele frequencies in our work is most likely the result of a recent expansion associated with the population recovery after the genetic bottleneck during the last glacial period. Genetic signals from the mid or early Pleistocene, when peas first expanded from the Near East westwards, were probably lost from recent populations due to the effect of other evolutionary forces. We detected significant clines in allele frequencies suggesting expansion routes from the West to the East (Cl3 → Cl6 & Cl1) and from Turkey to the South (Cl4 → Cl1; Cl5 → Cl2) and further to the East (Cl4 → Cl1; Cl5 → Cl2; Cl1 → Cl6; Fig. 5). It seems like population Cl4, which is located around the Black Sea expanded towards the South and East. At the same time the European cluster expanded eastwards. It is possible that the southward expansion of glaciers during the last ice age reduced the size of European populations of *P. s.* ssp. *elatius* and forced those into refugia as was observed in numerous Mediterranean plant species (Médail & Diadema, 2009). Following the retreat of glaciers, pea populations may have expanded from those refugia northward as well as eastwards. A process that could still be ongoing. Such an eastward expansion from refugia was also observed in other pan-Mediterranean legumes like narrow-leafed lupin (*Lupinus angustifolius*; Mousavi-Derazmahalleh et al., 2018) and pitch trefoil (*Bituminaria bituminosa*; García-Verdugo et al., 2021). The main difference between the *Lupinus* and *Bitumenaria* cases and wild peas is that the center of diversity of narrow-leafed lupin and pitch trefoil is located in the western rather than the eastern Mediterranean. Those species likely originated from the western Mediterranean, while the origin of *P. s.* ssp. *elatius* is located in eastern Mediterranean. The range expansion we detected may be the result of an eastward expansion from glacial refugia of the European populations which mixed with the eastern populations during the expansion and a simultaneous expansion of Turkish populations to the South and further to the east.

**Figure 5:**
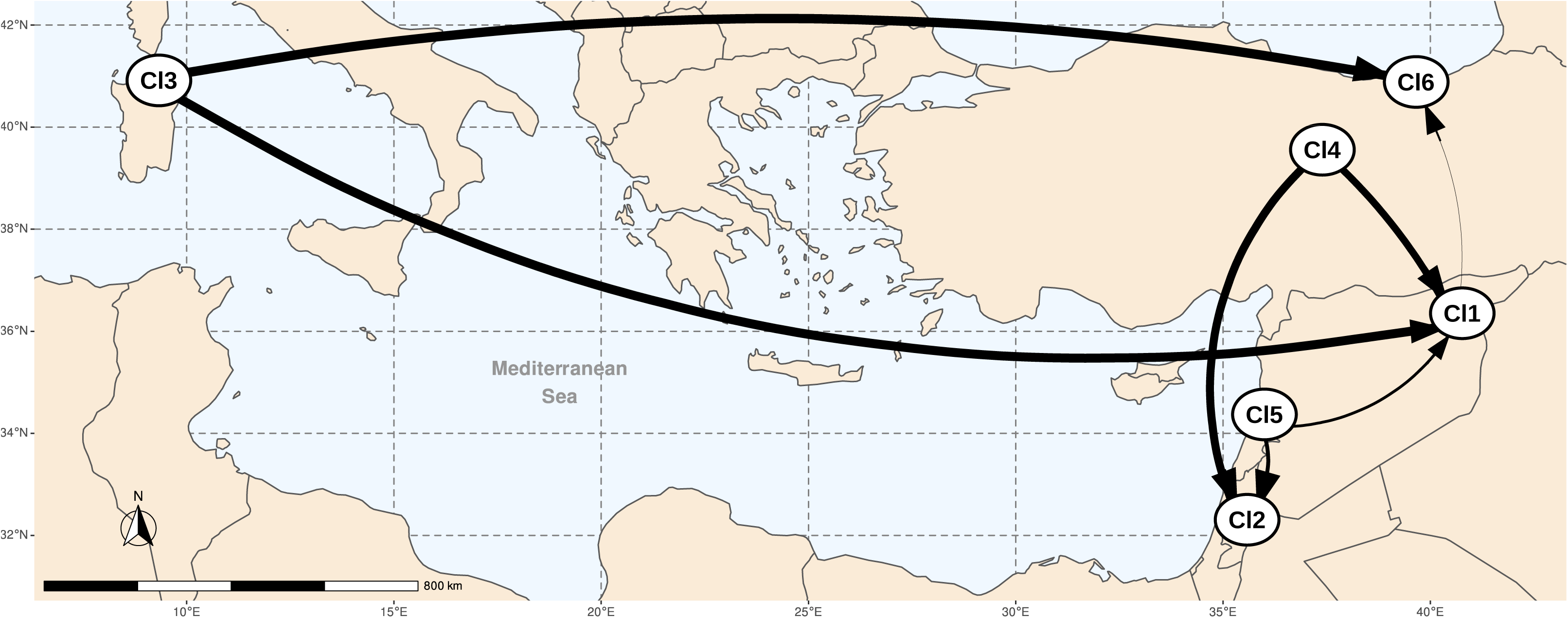
Distribution centers of genetic clusters with expansion routes estimated by pairwise directionality indices ψ. Arrows indicate the putative directions of expansions. Arrows’ thickness are proportional to the respective values of ψ. Only significant ψ are depicted (for exact values of ψ and their respective p-values see Tab. S5).

Overall, the interplay between environmental factors and recent evolutionary history affected the distribution of genetic diversity of *P. s.* ssp *elatius* where each subpopulations were differently affected by those factors and processes.

## Authors’ credit statement

**Timo Hellwig**: Conceptualization, Methodology, Investigation, Data curation, Formal analysis, Writing; **Shahal Abbo**: Review & Editing, Supervision, Funding aquisition; **Ron Ophir**: Review & Editing, Data curation, Supervision, Funding acquisition.

## Acknowledgements

We thank Dr. Naama Teboul for their critical comments on DNA extraction and Prof. Zvi Peleg for hosting T.H. in his lab during the preparation of DNA samples. The work was supported by The Israel Science Foundation [Grant/Award Number 307/17 to RO and SA]. Shahal Abbo is the incumbent of The Jacob & Rachel Liss Chair in Agronomy at the Hebrew University of Jerusalem. Timo Hellwig is a recipient of The Robert H. Smith stipend.

## Supplementary Figures

**Figure S1:**
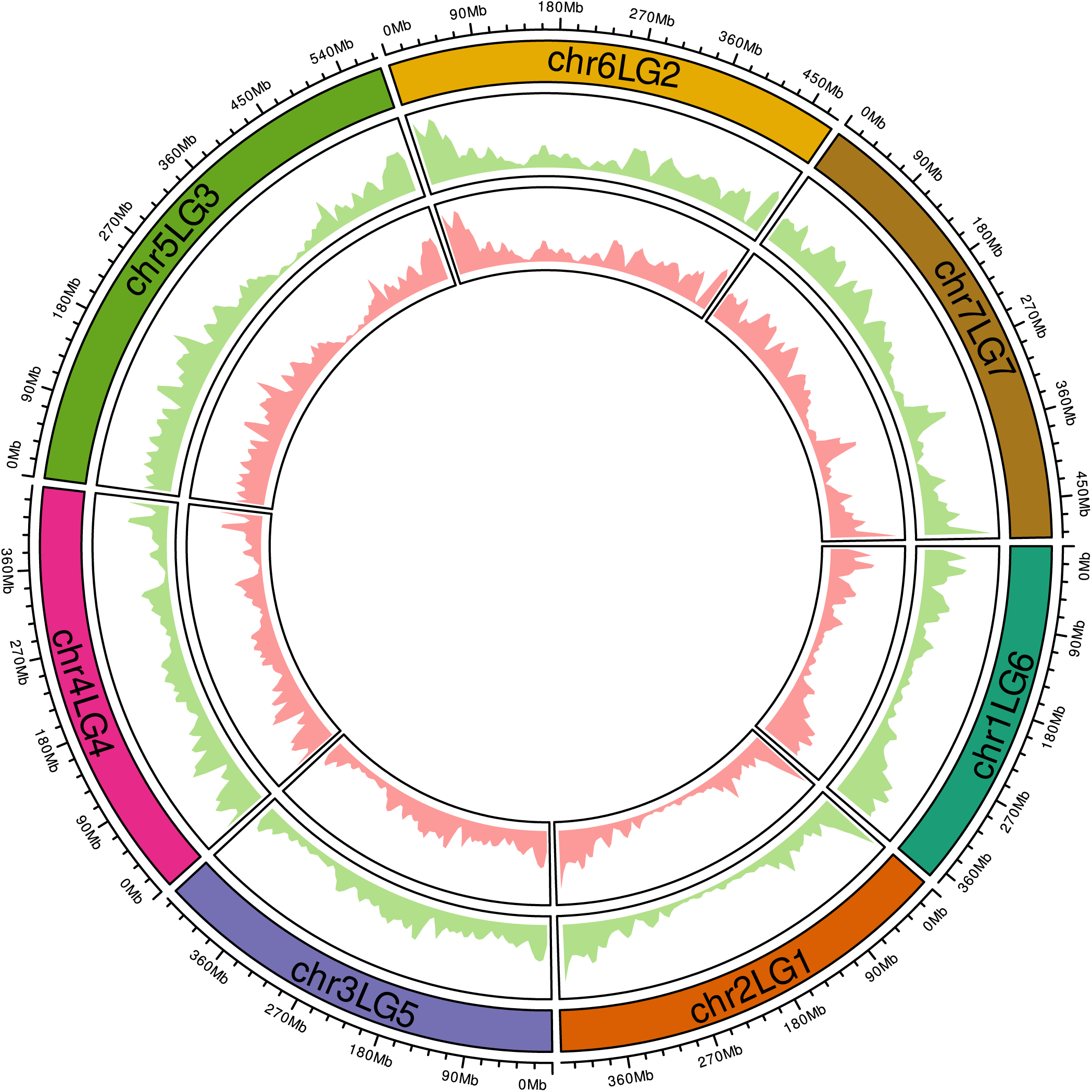
Distribution of SNP density over linkage groups. Outer circle (green): filtered variant set. Inner circle (red): LD pruned variant set.

**Figure S2:**
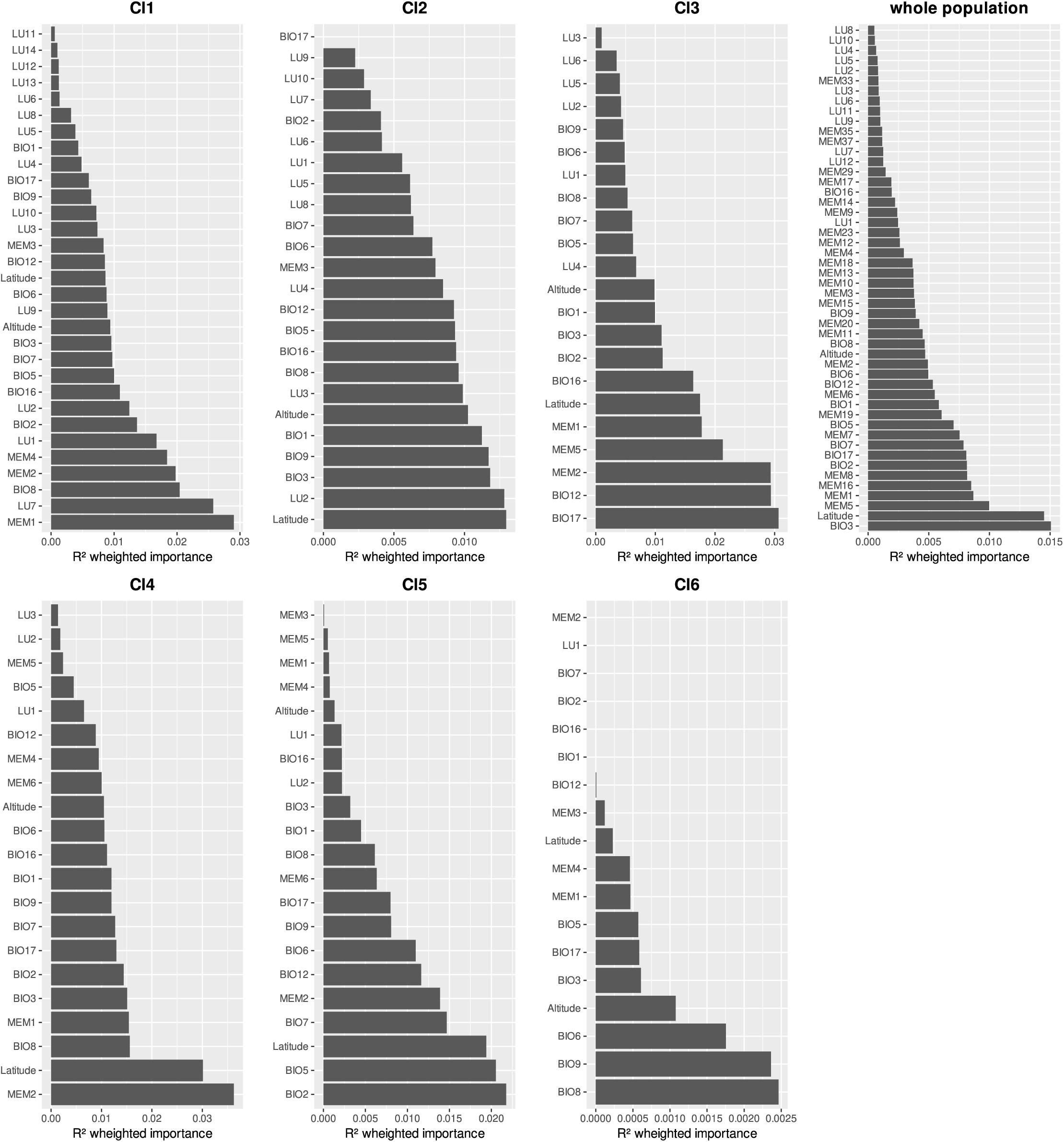
R²-weighted importance estimated in gradient forest analysis of each individual predictor for each genetic cluster and the whole population.

**Figure S3:**
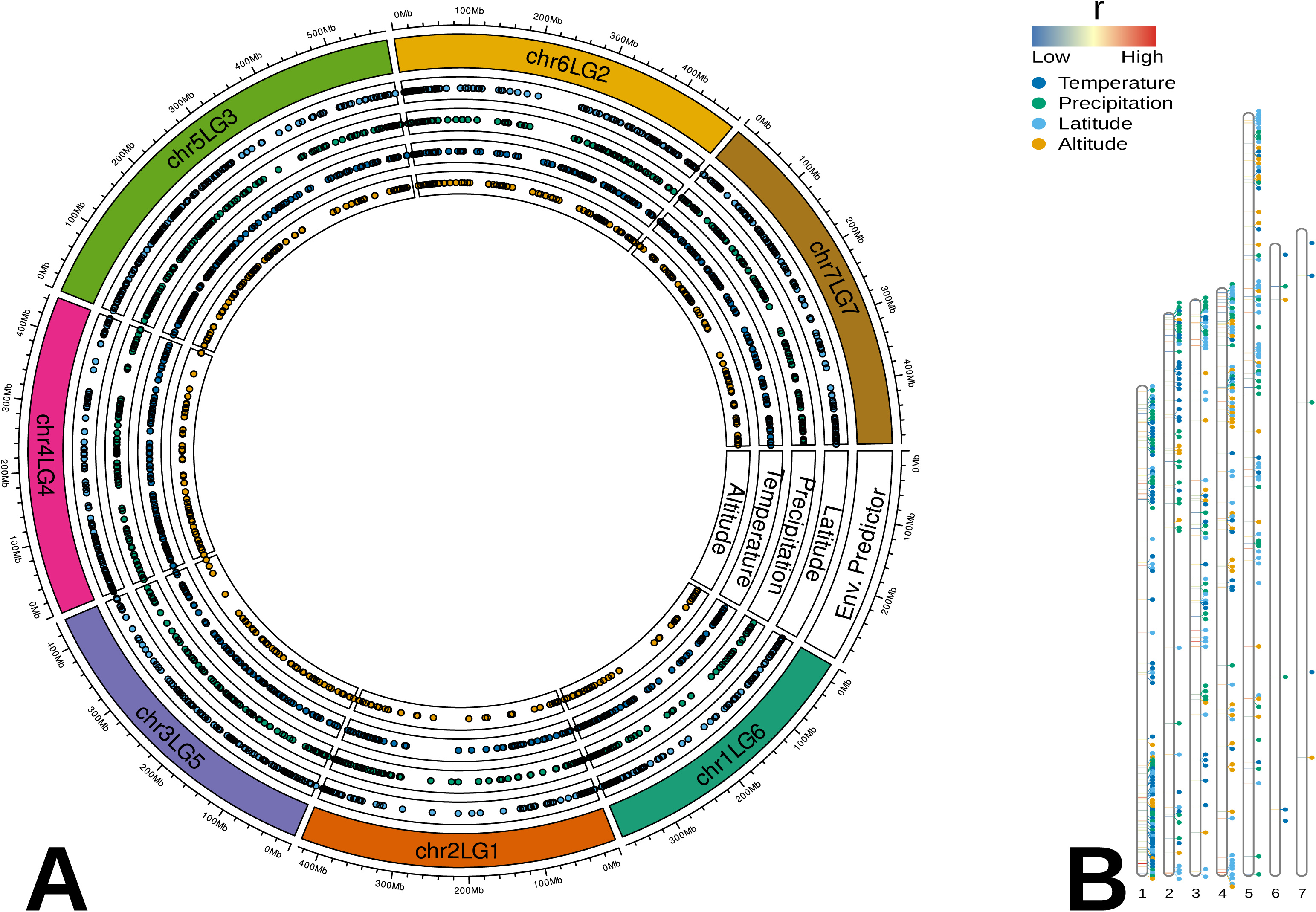
Genome-wide distribution of the SNPs associated with environmental factors. A: entire set of candidates. B: 10 % of candidates of each environmental predictor with the highest r values in RDA.

**Figure S4:**
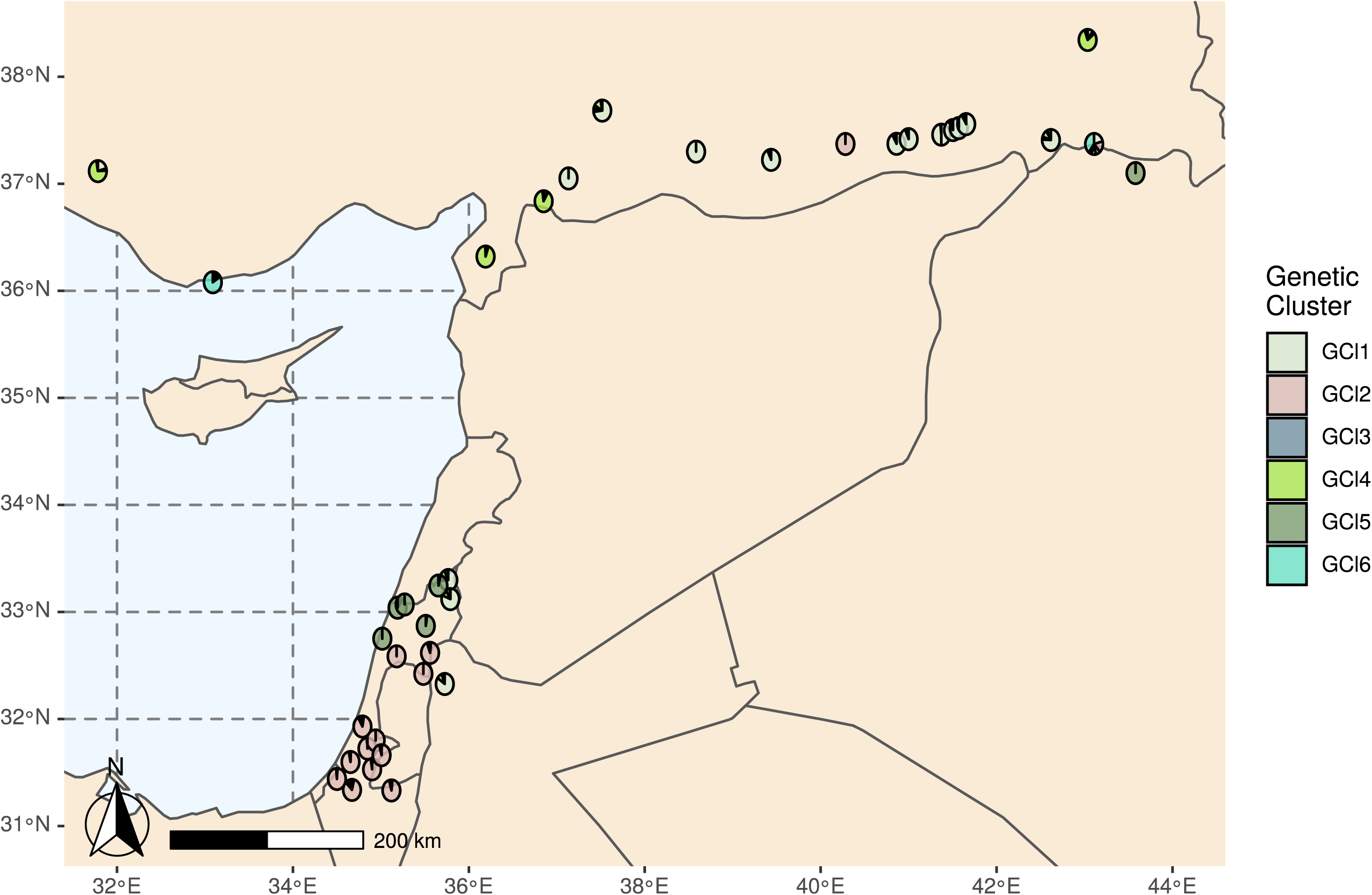
Detailed maps of Levantinian sample locations. Pie charts are located at the sample location of the analyzed accessions and represent sNMF subpopulation fractions with K=6 (for a map depicting the entire sample collection see Fig. 1).

